# Neural assemblies coordinated by cortical waves are associated with waking and hallucinatory brain states

**DOI:** 10.1101/2023.05.22.540656

**Authors:** Adeeti Aggarwal, Jennifer Luo, Helen Chung, Diego Contreras, Max B. Kelz, Alex Proekt

## Abstract

The relationship between sensory stimuli and perceptions is brain-state dependent: in wakefulness stimuli evoke perceptions; under anesthesia perceptions are abolished; during dreaming and in dissociated states, percepts are internally generated. Here, we exploit this state dependence to identify brain activity associated with internally generated or stimulus-evoked perception. In awake mice, visual stimuli phase reset spontaneous cortical waves to elicit 3-6 Hz feedback traveling waves. These stimulus-evoked waves traverse the cortex and entrain visual and parietal neurons. Under anesthesia and during ketamine-induced dissociation, visual stimuli do not disrupt spontaneous waves. Uniquely in the dissociated state, spontaneous waves traverse the cortex caudally and entrain visual and parietal neurons, akin to stimulus-evoked waves in wakefulness. Thus, coordinated neuronal assemblies orchestrated by traveling cortical waves emerge in states in which perception can manifest. The awake state is privileged in that this coordination is elicited by specifically by external visual stimuli.

## Introduction

Perceptual experiences are seamlessly integrated. Natural visual stimuli appear as meaningful scenes with an arrangement of objects, each one of which is itself a complex stimulus made up of multiple features, such as position, shape, texture, and color. The integration of the different features of the stimuli into objects happens subconsciously and is a *sine qua non* of our perceptual experiences. Neurons that encode different features of the visual world are distributed broadly among many cortical regions. Thus, to produce integrated perceptual experiences, neuronal activity distributed among many cortical areas must be appropriately coordinated. Zero phase lag synchrony of neural oscillations across different regions has long been hypothesized to play a role in mediating this coordination^1–6^. However, recent studies revealed that oscillatory brain signals exhibit a rich repertoire of phase relationships and resemble traveling waves that traverse the cortex in different directions and at different frequencies^7,8^.

Mounting evidence suggests traveling waves coordinate the neuronal activity essential for sensory perception. These waves are reliably evoked by stimuli^9–13^ and coordinate firing of individual neurons across different cortical regions^10^. Traveling wave patterns switch during spontaneously alternating perceptions observed during binocular rivalry^14^ and are associated with perceptual echoes^15^. Traveling waves can be used to implement predictive coding in naturalistic settings^16,17^. However, several observations complicate the relationship between traveling waves and perception. Traveling waves are observed in states where conscious perception occurs reliably, such as during normal wakefulness^9–11,18^, but also during sleep,^19,20^ and general anesthesia^18,21^. While traveling waves of activity are evoked by sensory stimuli^10,11,22^, they also occur spontaneously^22,23^. However, a detailed understanding of the essential differences between spontaneous and evoked traveling waves in different states of consciousness is lacking.

To address this knowledge gap, we use high density neurophysiological recordings to study the relationship between spontaneous and visual evoked traveling cortical waves in mice across three fundamentally distinct states of consciousness. During wakefulness, sensory perceptions are reliably evoked by suprathreshold stimuli. Under isoflurane anesthesia, subjects are unresponsive to all stimuli, exhibit slowing of cortical activity, and rarely report having perceptual experiences^24^. During the third state of ketamine-induced dissociative anesthesia, subjects are unresponsive to external stimuli but exhibit wake-like cortical levels of activity^25–29^ and report having vivid hallucinations^30–35^. While it is difficult to be certain about the specific timing of these hallucinations, the conjunction of behavioral reports and neurophysiological recordings suggests that these perceptual experiences likely occur during ketamine-induced unresponsiveness.

While investigations of neurophysiological underpinnings of hallucinations and dreams is of fundamental importance for the study of perception in general, these phenomena are extremely difficult to study, especially in animal models. Here, we chose to use ketamine to induce a dissociated state in mice based on several lines of evidence. Ketamine, at both anesthetic and sub-anesthetic doses, reliably induces sensory hallucinations in humans^30–35^. The neurophysiological activity patterns induced by ketamine in the mouse brain^36^ are similar to those induced by ketamine in humans^37^. In fact, a similar oscillatory neurophysiological activity pattern is observed in mice under ketamine and in humans who experience sensory hallucinations during the prodromal stage of seizures^36^. While most conventional sedatives and anesthetics induce brain activity resembling slow wave sleep^38^, activity induced by ketamine has some essential similarities to rapid eye movement (REM) sleep^39^—a stage of sleep typically associated with vivid dreaming^40^. Much like REM sleep, the ketamine-induced state is associated with decreased responsiveness to external stimuli^26,36^. Stereotypical behaviors exhibited by mice under ketamine share similarities to behavioral signatures of other hallucinogens in rodents^26^. Thus, while one can never directly test for the existence of hallucinations, or any other subjective experience^41^ in an animal model, the neurophysiological state induced by ketamine in mice exhibits essential similarities to states associated with dreaming and hallucinations in humans.

Based on these considerations, our primary goal was to characterize the relationship between spontaneous and stimulus evoked activity using a combination of high density ECoG and single unit recordings in visual and higher-order parietal cortices of mice during wakefulness, under isoflurane anesthesia, and in the ketamine-induced dissociative state. We show that visual stimuli reliably reset the phase of spontaneous 3-6 Hz cortical waves to give rise to feedback travelling cortical wave which orchestrates neuronal firing in the visual and parietal cortices. This phase reset is observed exclusively in the waking state. Interestingly, exclusively under ketamine, spontaneous 3-6 Hz cortical waves have several essential similarities to travelling waves elicited by stimuli during wakefulness. These spontaneous waves under ketamine also propagate in the feedback direction and coordinate neuronal firing across brain areas. Thus, an oscillatory neuronal assembly orchestrated by a feedback travelling cortical wave at 3-6 Hz is a promising neurophysiological mechanism for coordinating neuronal firing across the neocortex required for integrated sensory perceptions.

## Results

We performed high density *in vivo* electrophysiological recordings in head fixed mice (n = 32) that received visual stimuli consisting of either a 100ms full screen white flash of varying luminance (Methods) or a 10ms flash of a green LED (Figure 1a, Methods). A 64 channel electrocorticography (ECoG) grid placed over the dural surface of the contralateral (left) hemisphere was used to record local field potentials (LFPs) from the visual, association, retrosplenial, somatosensory, and motor/frontal areas. In a subset of animals (n = 14), two 32 channel laminar probes were also inserted perpendicularly into the cortex through holes in the ECoG grid to target the primary visual cortex (V1) and the posterior parietal area (PPA). In these animals, histological localization of the laminar probes was used to determine the stereotaxic locations of ECoG electrodes and enabled comparisons of activation patterns across mice (Methods).

**Figure 1:**
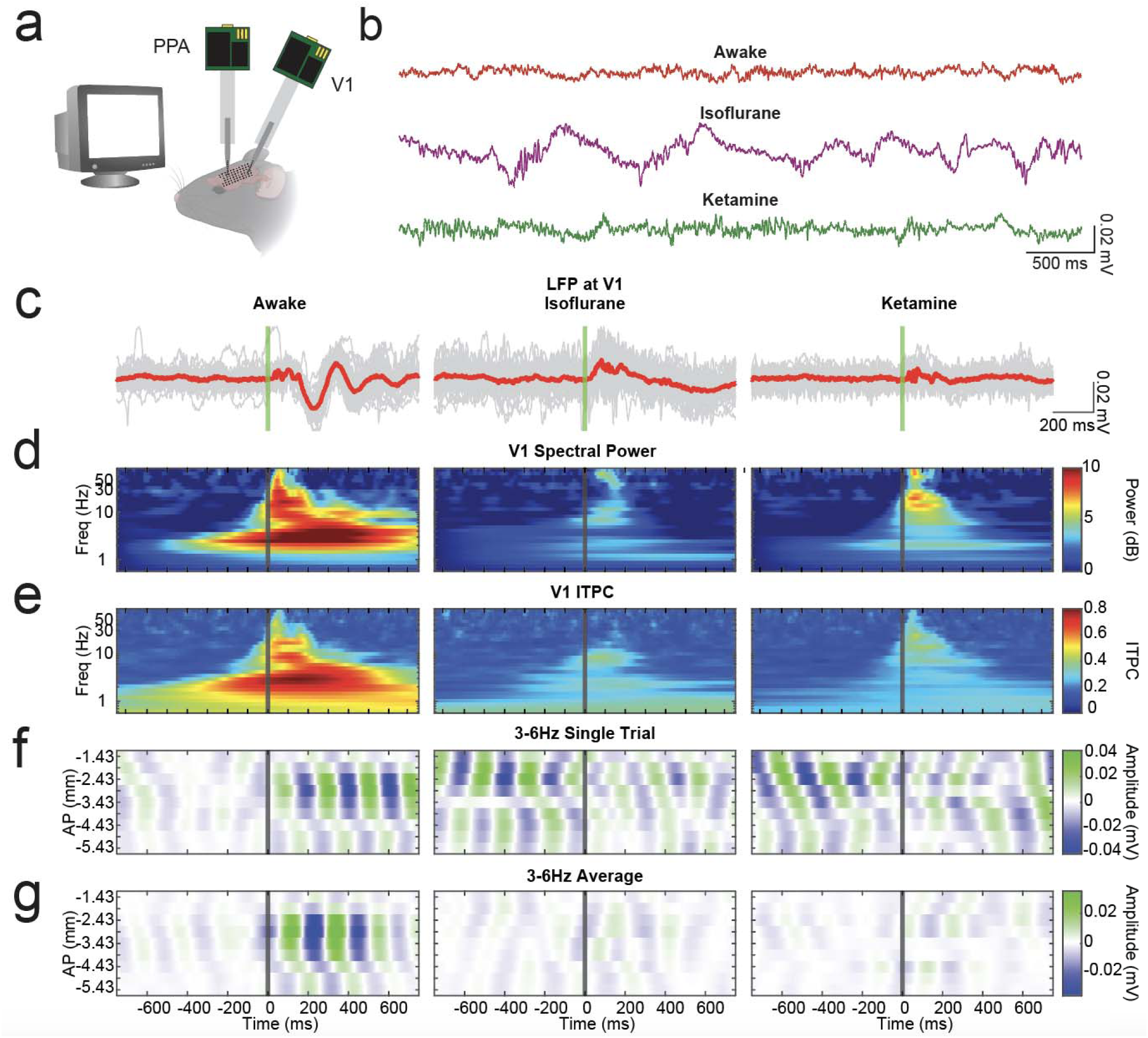
Spontaneous cortical waves exist during wakefulness, under isoflurane, and under ketamine, but are exclusively phase reset by visual stimuli in the awake brain. A. Experimental design: 64 channel electrocorticography (ECoG) grid used to record local field potentials (LFPs) from the cortical surface of the left hemisphere (n=32 mice). In addition, 32 channel laminar probes were placed in the primary visual cortex (V1) and the Posterior Parietal Area (PPA) in (n=14 mice). Stimuli consisted of 100ms of full screen flashes (44% brightness, 33 cd/m^2^) delivered by a CRT monitor placed in front of the right eye (Methods). B. Five seconds of spontaneous LFP recorded over V1 in the same mouse in the awake state (top), under isoflurane (middle), or under ketamine (bottom). C. 40 Single trials (gray) and average (red) visual evoked potentials (VEPs) from a representative mouse during wakefulness (left), under isoflurane (middle), and under ketamine (right). The vertical green line denotes stimulus onset. D. Stimulus evoked changes in spectral power (relative to pre-stimulus baseline) in V1 averaged over animals during wakefulness (left), under isoflurane (middle), or under ketamine (right). The vertical gray line denotes stimulus onset. E. Intertrial phase coherence in V1 averaged over animals during wakefulness (left), under isoflurane (middle), or under ketamine (right). The vertical gray line denotes stimulus onset. F. Single trial of LFPs filtered at 3-6Hz recorded in a column of electrodes (each row is a single channel) at −2.0mm lateral from bregma from the same representative mouse as in A. Note that spontaneous waves are observed in all brain states. In contrast to the awake state, under both isoflurane and ketamine, the phase of the slow wave is unaffected by the stimulus. G. Intertrial average VEP filtered at 3-6Hz from the same representative mouse as in A, F. Because the stimulus does not phase reset the spontaneous waves under isoflurane or ketamine, the waves observed in a single trial cancel out. In contrast, because the stimulus reliably resets the phase of spontaneous waves in the awake brain, a consistent feedback wave is observed after the stimulus.

To determine how the traveling waves of cortical activity evoked by simple visual stimuli^10^ are affected by the state of the brain, we recorded visual evoked activity in the same mice during wakefulness, under isoflurane, and under ketamine (Methods, Sup. Figure 1). Spontaneous cortical activity during these three states is shown in Figure 1b. Consistent with previous findings^26,42^, isoflurane increases low frequency oscillations, while brain activity under ketamine is similar to that observed in the awake state, albeit with moderately increased high frequency (gamma) activity (Figure 1b, Sup. Figure 1).

### 3-6Hz waves are present in spontaneous activity in all states, but visual evoked 3-6Hz waves are observed only in the waking state

Prior work^10^ has shown that the visual evoked potential (VEP) in mice can be decomposed into two oscillations: early gamma (30-50Hz) oscillations, and late 3-6Hz oscillations. Figure 1c shows that the early gamma oscillations are consistently observed within the first 100 ms of the stimulus in all three states. Consistent with this, gamma power (Figure 1d) in the first 100ms of the VEP is increased during wakefulness, under isoflurane, and under ketamine. Gamma ITPC induced by intermediate-intensity stimuli under isoflurane is reduced relative to wakefulness and ketamine. More intense stimuli, however, reliably evoked early coherent gamma under isoflurane (Sup. Figure 2). As we and others^10,43^ have shown, these gamma oscillations originate in the cortical input layer 4 and then propagate to supra- and infragranular layers. Consistent with this, the initial layer 4 sink supra-and infra-granular sources pattern is observed all three states (Sup. Figure 3a). Thus, in all studied states, the primary visual cortex continues to receive and respond to inputs from the thalamus (Sup. Figure 3).

**Figure 2:**
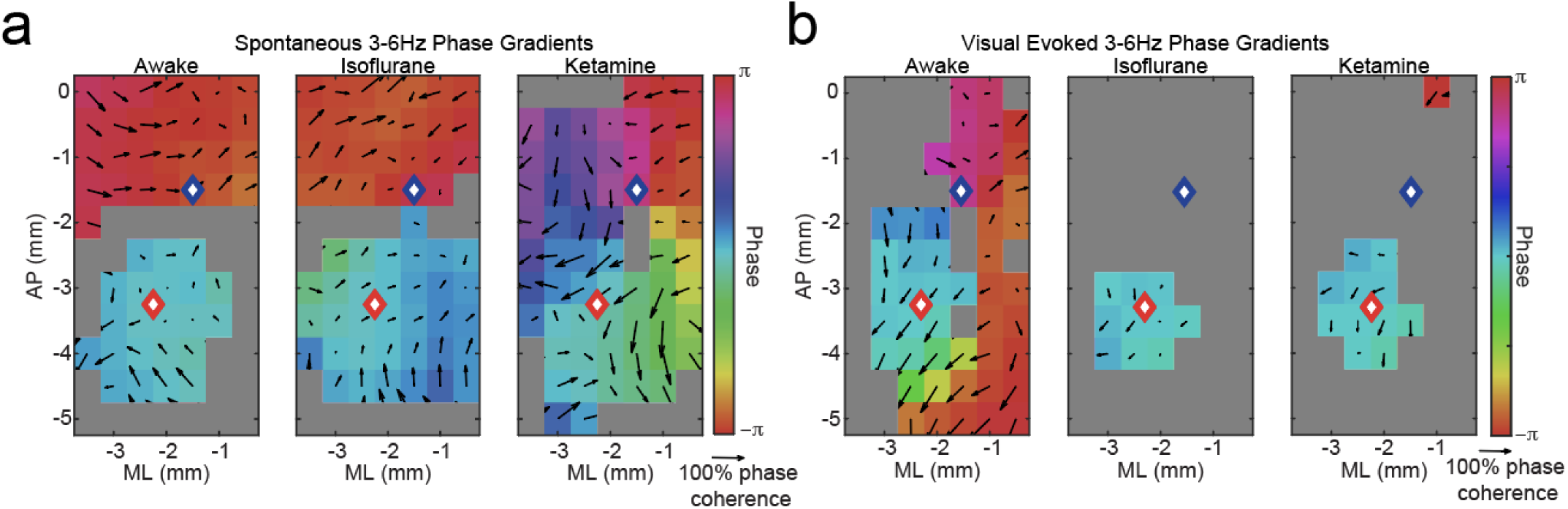
Spontaneous waves under ketamine and visual evoked waves in the awake brain both travel in the feedback direction. A. At each stereotaxic location, the phase averaged across animals (n=14) of the spatial component of the largest 3-6Hz spatiotemporal mode (Methods) observed during spontaneous activity is shown by color. All phases are computed relative to V1 (red diamond). PPA is shown by the blue diamond. The arrows show the spatial phase gradient averaged across trials and animals. The directions of the arrows show the direction of spatial phase gradient. The length of the arrows encodes phase coherence (Methods). Locations shown in gray did not meet statistical significance for phase coherence over trials computed using Raleigh test and adjusted for multiple comparisons using Bonferroni correction. B. Same analysis as in A performed on the spatial component of the most visually responsive 3-6Hz spatial mode (Methods). Note that visually responsive 3-6Hz waves during wakefulness reliably propagate in the feedback direction and involve much of the cortical surface. In contrast under both isoflurane and ketamine anesthesia, visual evoked waves are confined primarily to V1. Furthermore, note that spontaneous waves under ketamine and visual evoked waves in the awake state both exhibit caudal propagation patterns.

**Figure 3:**
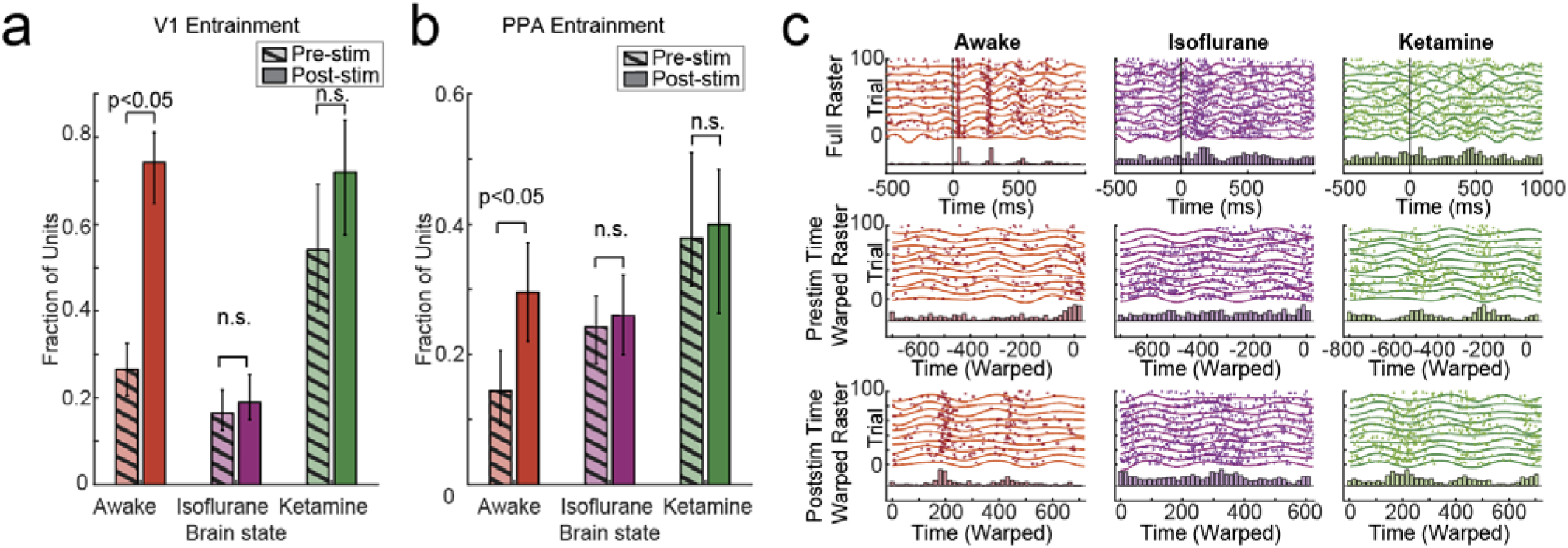
V1 and PPA neurons are entrained to the 3-6Hz visual evoked wave during wakefulness and to the spontaneous 3-6Hz under ketamine. A. Fraction of single units entrained to the slow 3-6Hz oscillation in V1 averaged across animals (n=11) in the awake state, as well as under isoflurane or ketamine anesthesia. Hashed bars show pre-stimulus, solid bars show post-stimulus. Error bars show standard error computed across animals. In the awake state, the fraction of entrained neurons is increased relative to the pre-stimulus baseline (χ^2^_V1_ = 185.9201, p_V1_ <10^-10^ Chi Square; p_AwakeV1_ <10^-10^, Tukey post hoc). Under ketamine (p_KetV1_ 0.435, Tukey post hoc) or isoflurane (p_IsoV1_ = 0.987, Tukey post hoc) no statistical differences in the number of entrained neurons are detected. Under ketamine, but not isoflurane anesthesia, the fraction of entrained neurons is comparable to that observed after the stimulus in the awake state (p_AwakeKetV1_ = 0.999, Tukey post hoc). B. Similar to A, but for cells in the Posterior Parietal Area (PPA). The number of PPA neurons entrained to the 3-6Hz wave increases after the stimulus presentation awake mice (χ^2^_PPA_= 25.9103, p_PPA_ = 9.2883 x 10^-6^, Chi-square; p_AwakePPA_ = 0.034, Tukey post hoc), but not under isoflurane (p_IsoPPA_ = 0.998, Tukey post hoc), or ketamine (p_KetPPA_ = 1, Tukey post hoc). The number of PPA neurons entrained by the spontaneous wave under ketamine is similar to that in the post-stimulus period in the awake state (p_AwakePPA_ = 0.034, Tukey post hoc). C. Example raster plots (top row) showing 100 trials (y-axis) from a representative V1 neuron firing at specific time points (x-axis) in relation to the visual stimulus (time t = 0 ms), in mice that are awake (left column), under isoflurane (middle column), or under ketamine (right column). Spike histogram is shown below the raster. The second row shows raster plots of the same neuron, after time warping to match the phases of the 3-6Hz oscillations in the pre-stimulus period (Methods). There is little entrainment before the stimulus when the mouse is awake or under isoflurane, but there is robust entrainment when the mouse is under ketamine. The third row shows a similar time-warped raster for the post-stimulus period 3-6 Hz. There is robust entrainment of cells to the slow wave when the animal is awake or under ketamine, but there remains little entrainment when the animal is under isoflurane.

In contrast to gamma oscillations, 3-6 Hz oscillations robustly increase in power and intertrial phase coherence after the stimulus in awake mice compared to mice under both ketamine and isoflurane (Figure 1d,e). Spontaneous LFP in all three brain states is dominated by low frequency activity. For example, LFP under isoflurane contain more low frequency power than during wakefulness (Figure 1b, Sup. Figure 1b). In contrast to the awake state, however, there is no further increase in 3-6Hz power or coherence evoked by the stimulus under ketamine or isoflurane (Figure 1d). Because the 3-6Hz oscillation is strongly affected by the state of the animal, we focus the rest of our analysis on the relationship between spontaneous and stimulus-evoked 3-6Hz activity across different states.

### Spontaneous and evoked 3-6Hz wave activity is organized into standing and traveling waves

As we have shown^10^, visual stimuli evoke 3-6Hz oscillations that propagate in the feedback direction from the higher order cortices towards V1. However, wave-like activity patterns in this frequency range are also observed spontaneously in all three states. For example, Figure 1f displays 750ms of pre- and post-stimulus LFP filtered at 3-6Hz across a column of electrodes along the anterior-posterior (AP) axis in a representative mouse. Note that in all three brain states, 3-6 Hz oscillations form coherent patterns of activity across space. In some instances, these patterns resemble a standing wave – oscillations are in opposite phases in the anterior and posterior aspects of the cortex and no systematic phase progression is observed (e.g. Figure 1f, isoflurane). In other cases, there is an orderly phase progression from anterior to posterior electrodes, and the overall pattern forms a traveling wave that propagates in the caudal direction (e.g. Figure 1f, ketamine).

During wakefulness, after the visual stimulus is presented, the 3-6Hz oscillations increase in power and are phase reset to organize into a feedback traveling wave. However, the onset of the visual stimulus fails to restructure the 3-6Hz activity when the same mouse is under isoflurane or ketamine. Consistent with this observation, the average of single trial LFPs filtered at 3-6Hz from the same mouse as in Figure 1f show a coherent visual evoked wave traveling posteriorly in the awake mouse (Figure 1g). In contrast, under both isoflurane and ketamine, the oscillations are not phase locked to the stimulus and cancel out, resulting in a greatly attenuated average signal. This suggests that while spontaneous waves in 3-6Hz are observed in all states, visual stimuli consistently reset the phase of these waves only in the awake brain.

### Spontaneous feedback waves under ketamine are similar to visual evoked feedback waves in awake mice

To characterize spontaneous and visual evoked waves across different animals and trials, we performed complex-valued singular value decomposition (SVD) separately on single trials of pre and post stimulus activity periods^10^. SVD factorizes the signals from the surface electrode grid into a set of mutually orthogonal spatiotemporal modes (Methods). To identify the most dominant spontaneous wave pattern, we examined the mode that captures the greatest fraction of the total signal. The first mode captures a similar fraction of variance in all three states (medianVariance_Awake_= 42.17%, IQR_Awake_ = 47.07%, medianVariance_Iso=_ 44.77%, IQR_Iso_ = 8.37%, medianVariance_Ket_= 45.17%, IQR_Ket_ = 48.54%). The spatial phase (color) and the spatial phase gradient (arrows) averaged across mice and trials in each of the three states is shown in Figure 2a. During wakefulness and under isoflurane, spontaneous oscillations in the anterior and posterior regions are in opposite phases and the phase gradients do not point in a consistent direction. Thus, on average, spontaneous activity during wakefulness and under isoflurane resembles a standing wave (Figure 2a). In contrast, in the same mice under ketamine, phase progression from anterior to posterior aspects of the cortex is clearly observed and the spatial gradients consistently point in the posterior direction. Thus, consistent with single trial data along a single column of electrodes (Figure 1f), spontaneous 3-6Hz oscillations under ketamine resemble a feedback traveling wave percolating along more than half of the cortical surface (Figure 2a).

Visual evoked waves were identified on a single trial level as the mode with the greatest increase in temporal amplitude during the post-stimulus period, relative to the pre-stimulus baseline^10^ (Methods). While in awake animals, the phases of the evoked waves are consistent across trials and mice^10^, in animals under isoflurane or ketamine, the intertrial and inter-animal variability dominates. Consequently, no consistent traveling wave is observed outside of V1 when animals are under isoflurane or ketamine (Figure 2b). This confirms our observation (Figure 1) that, in contrast to the awake state, visual stimuli under both ketamine and isoflurane do not consistently affect the phase of spontaneous cortical waves. Curiously though, spontaneous waves under ketamine and stimulus evoked waves in the awake state exhibit similar propagation patterns.

### Visual evoked 3-6Hz feedback waves in awake mice and spontaneous 3-6Hz waves in mice under ketamine entrain single unit neuronal firing in V1 and PPA

Low frequency LFP recorded from the cortical surface is thought to be dominated by post synaptic potentials in the superficial cortical layers^44^. Consistent with this view, we observe an oscillatory sink and source pattern in the superficial cortical layers of V1 at 3-6Hz range^10^ (Sup. Figure 3a). This raises the possibility that a wave of synaptic inputs percolating across the cortical surface may coordinate firing of neurons in different cortical areas. If this coordinated firing plays a role in perception, we would expect that 1) neuronal activity should transiently become coordinated by the 3-6Hz wave after the stimulus only in the awake state; 2) 3-6Hz waves should not entrain neurons under isoflurane either before or after the stimulus, and conversely 3) spontaneous waves should entrain neurons in mice under ketamine. Results in Figure 3 confirm all these predictions. In the awake state, the number of neurons entrained by the wave is significantly increased after the visual stimulus (Figure 3a,b, χ^2^ = 185.9201, p_V1_ <10^-10^, χ^2^ = 25.9103, p = 9.2883 x 10^-6^, Chisquare; p_AwakeV1_ <10^-10^, p_AwakePPA_ = 0.034, Tukey post hoc). In contrast, under both ketamine and isoflurane the number of entrained neurons is unaffected by the stimulus (Figure 3a,b, p_IsoV1_ = 0.987, p_IsoPPA_ = 0.998, p_KetV1_ 0.435, p_KetPPA_ = 1, Tukey post hoc). Furthermore, while the number of entrained neurons under isoflurane is consistently low both before and after the stimulus, the number of entrained neurons under ketamine is consistently high. Lastly the fraction of neurons entrained to the spontaneous 3-6Hz wave under ketamine is similar to that entrained by the stimulus evoked wave in the awake state (p_AwakeKetV1_ = 0.999, p_AwakeKetPPA_ = 0.578, Dunn-Sidak post hoc).

The raster plot in Figure 3c shows V1 neurons from the same representative mouse, reliably activated by the stimulus in all three states. In aggregate, the number of neurons that change their firing rate in response to the visual stimulus in V1 and PPA is similar in all three states (Sup. Figure 3c, Sup. Figure 4c). The key distinction between different states, thus appears to be the coordination of neural firing to the phase of the wave rather than the ability of the stimulus to alter the firing rate *per se*. The coordination between wave and firing activity is highlighted when time is warped (Methods) such that the phases of the 3-6Hz wave align (Figure 3c). In the awake brain, the firing is uncorrelated to the phase of the wave before the stimulus presentation, but becomes phase locked to the wave after the stimulus. In the animal under isoflurane, there is no association between the phase of the 3-6Hz wave and firing either before or after the stimulus. Under ketamine, in contrast, action potentials are phase locked to the 3-6Hz oscillation both before and after the stimulus. Thus, exclusively under ketamine, spontaneous waves entrain neuronal firing. Only in the awake state, stimulus evoked waves transiently entrain neuronal firing. This further underscores the essential physiological similarity between spontaneous activity under ketamine and stimulus evoked activity in the waking state.

**Figure 4:**
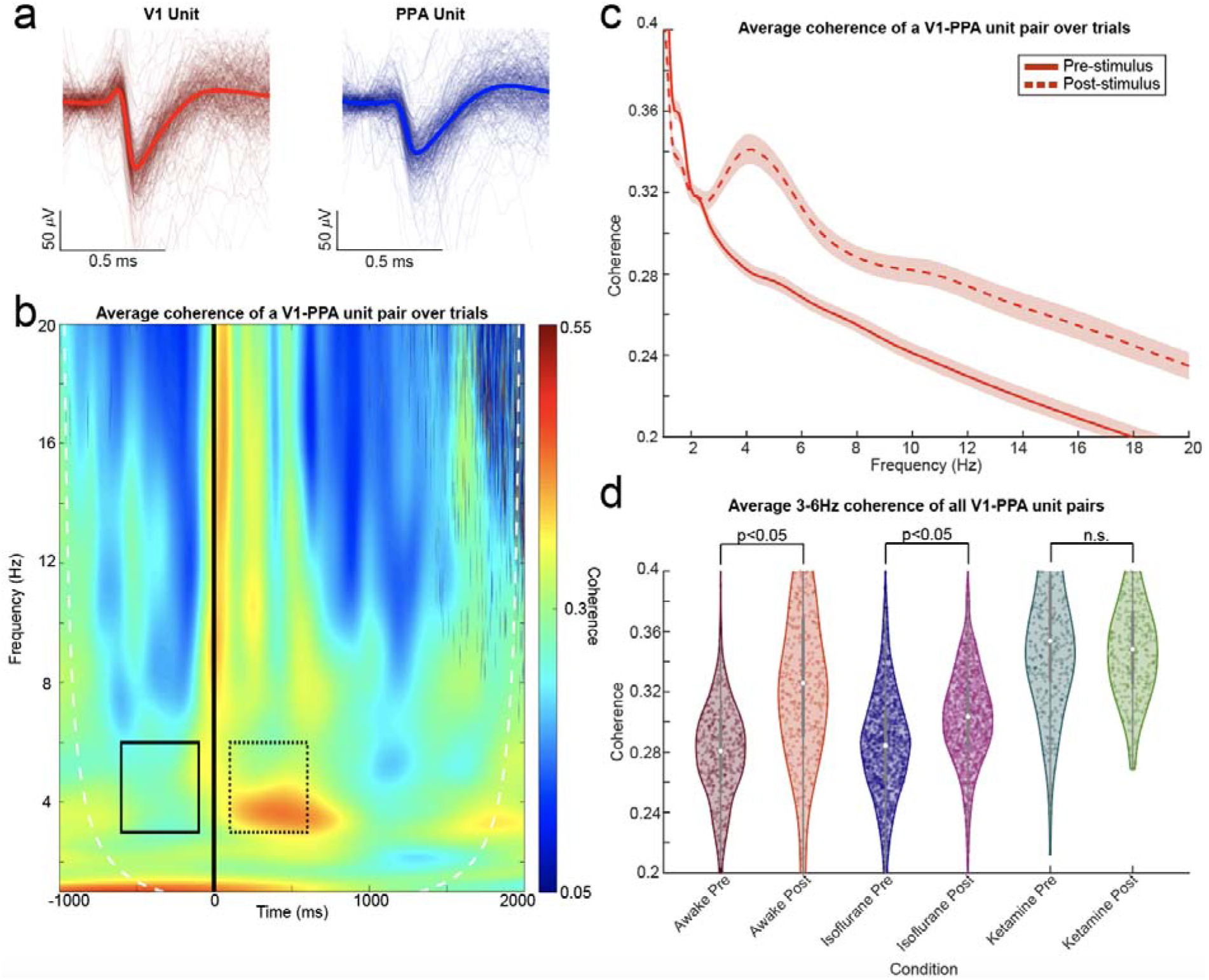
V1 and PPA cells form a transient oscillatory assembly at 3-6Hz after stimulus presentation in awake mice, and spontaneously under ketamine. A. Single-trial (thin lines) and average (thick lines) waveforms from representative V1 (left) and PPA (right) single units in a representative awake mouse. B. Average coherence of spike times from the V1 and PPA unit in A, averaged over trials. Time is on the x-axis with the flash occurring at 0ms (black vertical line). Frequency is on the y-axis, and coherence is represented by color. The solid black outline represents pre-stimulus time from - 600ms to −100ms before the flash onset. The dashed black outline represents post-stimulus time from 100ms to 600ms after the flash onset. The white dashed line represents the cone of influence, outside of which the coherence measure is inaccurate. C. Average coherence of all V1-PPA unit pairs over trials and animals in the −600ms to −100ms pre-stimulus (solid) and 100ms to 600ms post-stimulus (dashed) time frames in awake mice. After stimulus presentation, coherence between V1 and PPA neurons exhibits a peak at 3-6 Hz. D. Average 3-6Hz coherence of V1-PPA unit pairs over trials in each condition (p =1.1959 x 10^-172^, Kruskal Wallis; p_Awakepre-post_ < 10^-10^; p_Isopre-post_ < 10^-10^, p_Ketpre-post_ = 0.9999, p_KetPreAwakePost_ = 0. 0668, p_KetPreAwakePost_ = 0. 3514, p_KetPreAwakePost_ = 0. 0668, p_KetPreAwakePost_ = 0. 3514, Dunn-Sidak post hoc).

### Stimulus evoked 3-6Hz traveling waves orchestrate oscillatory neuronal assemblies spanning the visual and the parietal cortex

We hypothesized that because both V1 and PPA neurons become entrained to the visual evoked traveling wave in the awake brain, their firing ought to form a transient oscillatory assembly in the immediate aftermath of the stimulus. To test for the emergence of an oscillatory cell assembly specifically, we analyzed neuron correlations in the frequency domain using coherency. An example coherogram averaged across 100 trials of visual stimuli for a representative V1-PPA neuron pair recorded in the awake state (Figure 4a) is shown in Figure 4b. As expected, prior to the stimulus, the firing of V1 and PPA neurons is essentially uncorrelated in any frequency range. Roughly 200 ms after the stimulus, the activity exhibits a significant peak in coherence at 3-6Hz. This coherent activity lasts approximately 750 ms after which the correlated firing of V1 and PPA neurons decays and becomes uncorrelated once again. To characterize the robustness of this finding, we computed coherence as a function of frequency in the pre-stimulus and post-stimulus periods (−600ms to - 100ms, 100ms to 600ms respectively) for all 218 V1-PPA neuron pairs recorded in 10 mice (Figure 4c). This average coherence exhibits a clear peak confined to the 3-6Hz range (shading shows 95% confidence bounds). Thus, the correlations in firing between V1 and PPA neurons form an oscillatory assembly spanning V1 and PPA in the immediate aftermath of the stimulus in the awake brain.

To determine whether this oscillatory assembly emerges exclusively in the awake state after the stimulus, we computed coherence averaged across pre- and post-stimulus regions in the time-frequency plane (solid and dashed rectangle respectively, in Figure 4a) in all three conditions. Figure 4d shows this analysis pooled across (218 pairs from awake mice, 1946 pairs from mice under isoflurane, 208 pairs from mice under ketamine). Consistent with the results of Sup. Figure 5, V1 and PPA neurons under isoflurane exhibit less coherence both before and after the stimulus compared to neuron pairs in awake mice during the post-stimulus period (p_IsoPreAwakePost_ = < 10^-10^, p_IsoPreAwakePost_ = < 10^-10^, Dunn-Sidak post hoc). In contrast, the coherence between V1 and PPA neurons under ketamine is unaffected by the timing of the stimulus and is statistically indistinguishable from the coherence observed in awake post-stimulus neuron pairs (p_KetPreAwakePost_ = 0. 0668, p_KetPreAwakePost_ = 0. 3514, Dunn-Sidak post hoc). Thus, visual stimuli presented to awake animals elicit a transient formation of a 3-6Hz oscillatory activity pattern that is reflected both in the local field potentials and correlated firing of neurons across distant cortical regions. Under ketamine, in contrast, a similar oscillatory assembly is observed spontaneously and is unaltered by the stimulus. Finally, under isoflurane, no such assembly is formed either before or after the stimulus.

**Figure 5:**
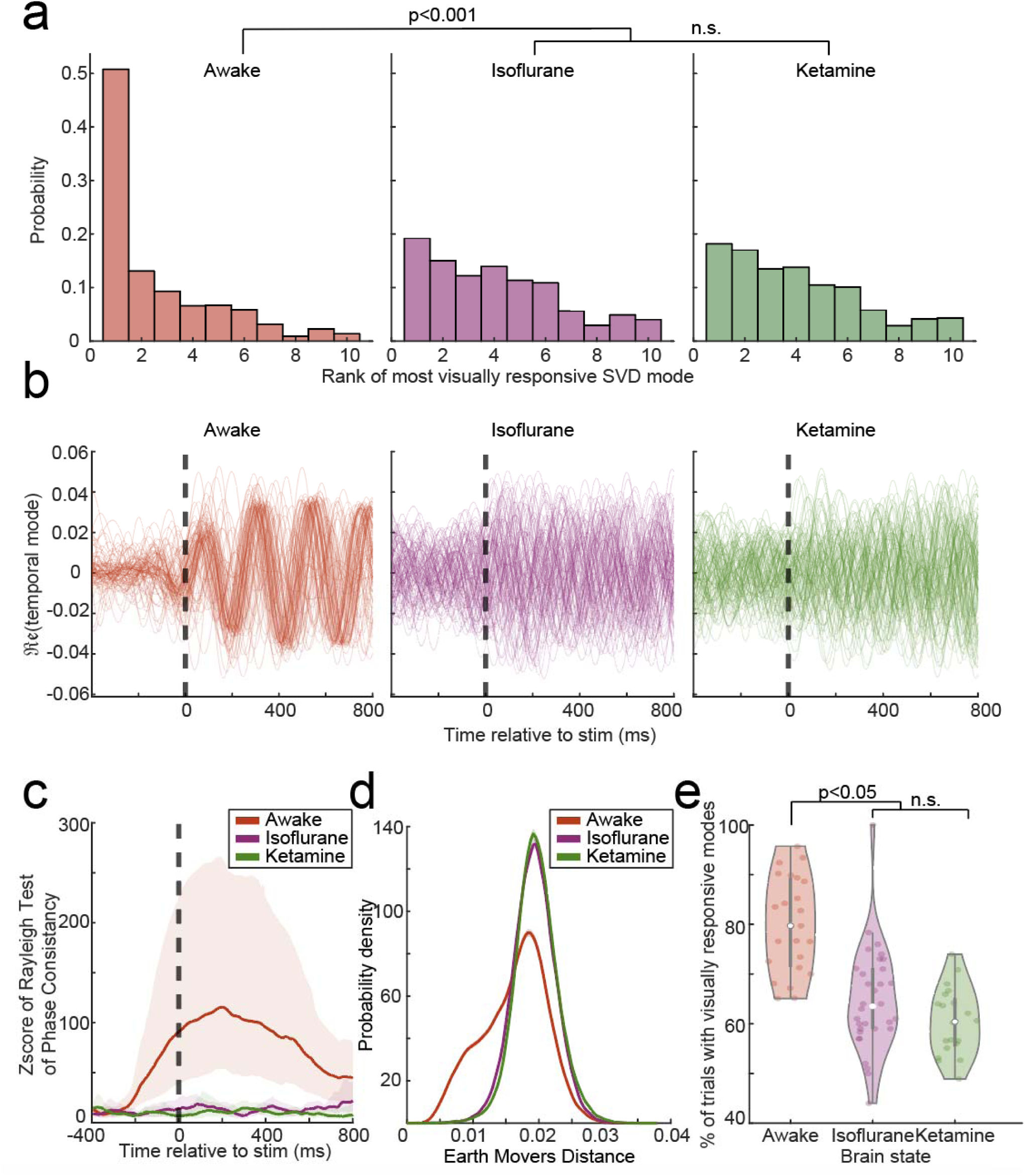
Visual evoked 3-6 Hz waves have high signal to noise ratio, consistent phase and are reliably elicited only in the awake mice. A. Distribution of ranks of the most visually responsive modes in each behavioral state. Visually responsive modes were more likely to be of lower rank (account for greater fraction of the signal) in awake mice than mice under isoflurane or ketamine (p =1.5655 x 10^-130^, Kruskal Wallis; p_Awake-Iso_ < 10^-10^; p_Awake-Ket_ < 10^-10^, p_Iso-Ket_ = 0.8531, Dunn-Sidak post hoc). B. Real part of the temporal component of the most visually responsive mode in a representative animal in the awake (right), isoflurane (middle) or ketamine (left) state. Each trace shows a single trial. The temporal components across trials align transiently after the stimulus in the awake animal, but not under ketamine or isoflurane. C. Deviations of phases of the most visually responsive mode from uniform circular distribution averaged across trials and animals expressed as Z score. Shading represents 95% confidence intervals bootstrapped over trials. The phases of the visually responsive mode deviate strongly from uniform distribution during wakefulness after stimulus presentation. This phase coherence decays to baseline by ∼ 800 ms. Under ketamine and isoflurane, no appreciable deviation from uniform distribution is observed after the stimulus. D. Distribution of Mean Earth Movers distance (EMD) between spatial amplitudes of the most visually responsive modes across trials and animals in the awake state (red), under isoflurane (purple), or under ketamine (green). Shading represents 95% confidence intervals. In the awake state, the spatial distribution of the most visually responsive mode is significantly more similar from trial to trial than under ketamine or isoflurane. No appreciable differences between ketamine and isoflurane are observed. E. Probability that a visually responsive mode was detected is higher in the awake state and under isoflurane or ketamine shown as a violin plot (each point is a probability estimated across all trials in a single animal) (p =3.966 x 10^-7^, Kruskal Wallis; p_Awake-Iso_ = 1. 1.6435 x 10^-6^; p_Awake-Ket_ = 2.4853 x 10^-5^, p_Iso-Ket_ = 0.9999, Dunn-Sidak post hoc).

### Characteristics of visual evoked 3-6Hz feedback waves in the awake brain indicate their potential neurophysiological significance in sensory perception

If visual evoked 3-6Hz waves were important for visual perception, these waves should have a high signal to noise ratio, should be consistently evoked on most trials, and should be phase locked to the stimulus. To determine if visual evoked 3-6Hz waves have high signal to noise ratio, we quantified the fraction of trials in which the most visually evoked mode accounted for the largest total fraction of variance. Indeed, we find that that the most visually responsive 3-6Hz wave tends to be the first SVD ranked mode in awake mice as compared to mice under isoflurane or ketamine (p =1.5655 x 10^-130^, Kruskal Wallis; p_Awake-Iso_ < 10^-10^; p_Awake-Ket_ < 10^-10^, Dunn-Sidak post hoc) (Figure 5a). We failed to detect any statistically significant differences in the signal to noise ratio between isoflurane and ketamine (p_Iso-Ket_ = 0.8531, Dunn-Sidak post hoc).

In awake mice, the stimulus resets the phase of spontaneous 3-6Hz oscillations (Figure 1f) and the phase of visual evoked 3-6Hz oscillations align over trials (Figure 1g, Figure 5b,c), whereas the distribution of the temporal phases of visual evoked waves across trials in mice under isoflurane or ketamine is more uniform (Figure 5b,c). To determine if the spatial activation profile of visual evoked slow waves was consistent across trials, we calculated the distribution of the Earth Movers Distance (EMD) (Figure 5d) and cosine distance (Sup. Figure 6) between the spatial amplitude of the most visually responsive mode between trials. We find the spatial activation pattern of visual evoked 3-6Hz waves are more consistent on a trial-by-trial basis than those found in mice under isoflurane or ketamine. Finally, to test for reliability, we quantified the number of trials in which at least one visually evoked mode (a mode in which the post-stimulus temporal amplitude exceeds pre-stimulus temporal amplitude by at least 6 standard deviations). We found that visual evoked 3-6Hz waves are more reliably elicited when animals are awake compared to when they are under isoflurane or ketamine (Figure 5e, p =3.966 x 10^-7^, Kruskal Wallis; p_Awake-Iso_ = 1. 1.6435 x 10^-6^; p_Awake-Ket_ = 2.4853 x 10^-5^, Dunn-Sidak post hoc). Again, we failed to detect any significant differences in reliability between ketamine and isoflurane states (p_Iso-Ket_ = 0.9999, Dunn-Sidak post hoc). Thus, visually evoked traveling waves account for much of the total signal and are reliably evoked by visual stimuli in a reproducible spatiotemporal pattern exclusively in the awake state.

**Figure 6:**
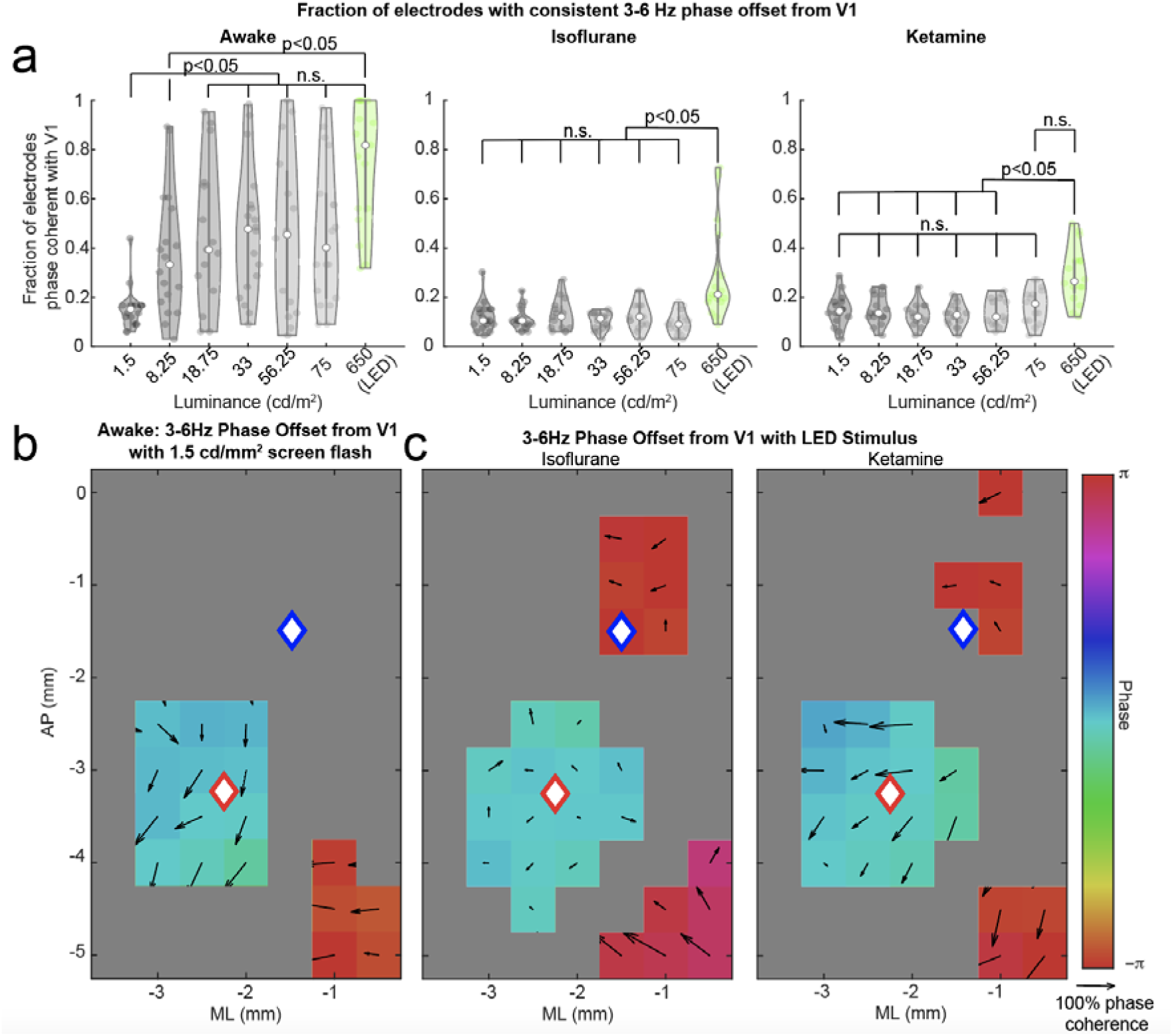
Visually evoked wave under isoflurane and ketamine resembles the wave evoked by subthreshold stimuli in awake mice. A. Fraction of electrodes in which phase of the slow waves is coherent with V1 (y axis) across trials within each mouse (individual points), when animals are shown 100ms screen flashes of varying intensities or an LED flash, during wakefulness (left), under isoflurane (middle) or ketamine (right). During wakefulness, the number of coherent electrodes increases in a stepwise fashion between 1.5 and 8.25 cd/m^2^ and remains approximately constant for higher intensity stimuli. Under both ketamine and isoflurane, the number of coherent electrodes is consistently low for stimuli of all intensity. B. At each stereotaxic location, the average phase offset of the most visually evoked mode from V1 (the red diamond) is plotted in color for lowest intensity (1.5 cd/^2^) stimulus recorded in awake mice. The arrows depict the spatial gradient. The directions of the arrows show the direction of spatial phase progression and the magnitude of the arrows correspond to the consistency of the angle of the spatial phase gradient over trials and animals. Gray locations did not meet Bonferroni corrected statistical significance (Raleigh test) across trials and animals. The blue diamond denotes the Posterior Parietal Area. C. Similar phase plots for maximum intensity stimuli (640 cd/mm^2^) recorded in mice under isoflurane and ketamine. In all three cases the visually evoked mode is confined to V1. Phases of the wave elicited by weak stimuli in awake mice resemble those obtained for high intensity stimuli under isoflurane and ketamine. Note that the phase plots and phase gradients are similar when shown the brighter LED stimulus to when mice are shown the dimmer screen flashes (see Figure 2). Awake data (left panel in B) is adapted from data presented in (Aggarwal et al, 2022), shown in Sup.Figure 8b) and is shown here for comparison to responses observed under ketamine and isoflurane.

### Visual Evoked 3-6 Hz feedback waves emerge in a stepwise function in awake mice, but fail to be evoked even at maximum intensity stimuli in mice under isoflurane or ketamine

Previously we have shown that as the intensity of the visual stimulus is increased, there is a sharp increase in the spatial phase consistency of visual evoked of 3-6 Hz feedback waves in the awake state^10^. Further, the sharp increase in the spatial extent of the wave occurs near murine perceptual threshold^45–47^. After the lowest intensity screen flash at 1.5 cd/mm^2^, only 20% of electrodes exhibit a consistent phase offset from V1 in the awake state (Figure 6a). When the stimulus intensity is increased from 18.25 cd/mm^2^ to 650 cd/mm^2^, the fraction of all electrodes phase locked to V1 is approximately double that for the weak stimulus and remains largely insensitive to stimulus intensity (Figure 6a). Remarkably, 3-6Hz waves elicited by maximum intensity screen or even LED flashes in mice under isoflurane or ketamine resemble the curtailed pattern of 3-6Hz waves mostly confined to V1. This pattern is essentially similar to that evoked by the weakest stimuli in awake mice (Figure 6 b,c). This similarity between responses to the weakest stimuli in the awake mice and otherwise strong stimuli in mice rendered unresponsive with either ketamine or isoflurane lends further evidence that supports the conclusion that perception of visual stimuli requires integration of activity across different cortical sites orchestrated by the 3-6Hz wave.

## Discussion

In our previous work^10^, we determined that in the waking brain, large scale 3-6Hz feedback traveling waves are evoked by visual stimuli. These results establish a correlation between traveling waves and visual perception. Building upon these previous findings, we pharmacologically manipulate the brain’s ability to perceive internal or external stimuli to determine whether properties of these waves depend on the state of the brain. We show that while 3-6 Hz traveling waves are observed spontaneously in awake mice, they do not entrain neuronal activity in V1 or PPA. Visual stimuli reliably reset the phase of spontaneous 3-6Hz oscillations to form a visual evoked traveling wave propagating caudally and correlate the firing of neurons in V1 and PPA specifically at 3-6Hz. Under anesthesia or in the ketamine-induced dissociative state, however, visual stimuli do not reset the phase of spontaneous waves at 3-6 Hz. Under anesthesia, V1 and PPA neurons are not correlated to the wave or to each other spontaneously or after stimulus presentation. In contrast, in the dissociative state in the absence of any stimulus, spontaneous waves resemble the awake brain’s stimulus-evoked waves both in terms of the propagation direction and the ability to correlate V1 and PPA neuronal firing. In summary, we show that a hallmark of sensory responsiveness is the ability of a stimulus to perturb spontaneous traveling waves. The capacity for sensory perceptions, be they stimulus evoked or hallucinatory, on the other hand, may be related to the formation of the 3-6Hz oscillatory assembly of neurons across the visual hierarchy.

The differences in the responses evoked by simple visual stimuli in different states of consciousness offer insights into the fundamental neurophysiological distinctions between them. Consistent with previous work^48^, we find that the early component of the evoked potential is preserved in all states (Figure 1, Sup. Figure 2). This early component of the evoked response, associated with gamma oscillations, reflects thalamic input into V1^43^. Indeed, we find that the early pattern of sinks and sources in V1 is similar among the three states (Sup. Figure 3). Further, approximately the same number of V1 neurons alter their firing rate in response to the stimulus in all three states. Thus, in the state of diminished responsiveness, the transmission of thalamic sensory inputs into the cortex is not dramatically disrupted. Similar observations have been made with auditory stimuli during slow wave sleep^49,50^ and under anesthesia^51^. We also observe that PPA—a higher order cortical region^52–54^ – continues to receive inputs in all states and approximately the same fraction of PPA neurons alter their firing rate in response to visual stimuli (Sup. Figure 4). Thus, the failure of signal propagation from the primary to higher order cortical areas does not readily account for the differences in the states of consciousness. The key distinguishing feature that appears to be unique to normal wakefulness is that visual stimuli evoke a coordinated oscillatory assembly^55,56^ of neurons orchestrated by the feedback traveling wave at 3-6Hz.

There is an extensive body of literature linking correlated oscillations in brain signals to perception^9,10,22^, attention^57^, cognitive control^58^, representation of space^59^ etc. While the existence of traveling waves is not in question, whether these correlated oscillations play a functional role or are epiphenomenal remains a hotly debated issue. One of the major reasons for this confusion is that oscillations *per se* and correlations among them can arise generically in a broad class of systems^8^. Thus, observing traveling waves does not directly point to their underlying mechanisms or behavioral significance. The waves that we and others^9,13,19,22, 60–66^ observe in the LFPs reflect the spatial patterns of synaptic potentials. The effect of this synaptic input, however, depends strongly on the specific network mechanisms that produce these synaptic inputs and the states of individual neurons that receive it.

The specific neuronal and circuit mechanisms were not the focus of this work, and their detailed understanding awaits future investigations. Nevertheless, recent computational modeling efforts^67^ suggest a general class of mechanisms that may explain our experimental results. Networks with predominantly local connectivity and conduction delays naturally produce traveling waves over a broad range of parameters^67^. In the regime where coupling between neurons is weak, these spontaneous traveling waves do not strongly entrain firing. Thus, there is no inherent contradiction between traveling waves of synaptic activity and asynchronous irregular and weakly correlated neuronal firing^68^. This regime is reminiscent of spontaneous brain activity during wakefulness observed herein and in previous studies^22^.

When the coupling strength between neurons is increased, neuronal firing becomes correlated and entrained to the phase of the traveling wave. Interestingly, as we demonstrate, only during wakefulness can a suprathreshold stimulus reliably switch the system from a weakly coupled to a more strongly coupled regime. The fact that neurons are spontaneously entrained to the wave and correlated to each other under ketamine, may result from elevated baseline coupling between cortical neurons that ketamine might induce. One potential mechanism that can give rise to this increased coupling is the preferential suppression of inhibitory cortical interneurons by ketamine^26,69^ and the consequent disinhibition of pyramidal cells^70^. Conversely, isoflurane is thought to disrupt cortico-cortical synapses^71^ and preferentially depress excitatory neurotransmission^72^. This decrease in coupling may explain why under isoflurane cortical neurons are never entrained by the wave. Interestingly, while ketamine preserves the overall level of cortical activity, the population of spontaneously active pyramidal neurons switches^26^. Thus, while the overall excitability of the cortical neurons may explain why under some circumstances the wave is able entrain neuronal firing, the identity of specific neurons that are recruited into the oscillatory assembly will depend strongly on the details of cortical microarchitecture.

The idea that specific oscillatory LFP dynamics may accompany sensory dissociation has been previously suggested. Both ketamine administration in mice and spontaneously occurring hallucinations in patients suffering from epilepsy are associated with coherent 1-3 Hz oscillations in brain activity^36^. Oscillations in the 3 Hz range were also identified under ketamine in patients implanted with ECoG electrodes for epilepsy localization^37^. It is clearly impossible to know whether mice experience hallucinations. However, the fact that LFP oscillations in humans during both ketamine-induced and spontaneously occurring hallucinations resemble those observed here suggests that at the neurophysiological level the state that we observe here in mice under ketamine is similar to that associated with self-reported hallucinations in humans. Previously it was unclear why a specific oscillatory behavior should give rise to sensory dissociation. Here, we demonstrate that waves of activity evoked by suprathreshold stimuli in the waking mouse brain are similar to those observed spontaneously under ketamine. This provides a link between specific cortical waves at 3-6Hz induced by ketamine and hallucinatory experiences unrelated to the environment.

We did not train mice to report when they see a stimulus and thus, the relationship between traveling waves and behavioral report will have to be elucidated in future work. Distilling which specific aspects of neuronal signals are associated with perception is a challenging task because brain activity associated with perception, reward anticipation, and movement become difficult to disentangle, especially on a single trial basis^73–76^. In altered states of consciousness, the behavior is further confounded by disruption of memory, motor coordination, and motivation.

Nevertheless, our results have important implications for the role of traveling cortical waves in perception. Classical theories assumed that the visual system processes stimuli in a hierarchical feedforward fashion^77,78^. In contrast, recent work suggests that while feedforward processing does indeed occur, the overall activity is heavily influenced by feedback^79^. In this view, the stimulus does not elicit a perception *per se*, but serves to modulate the spontaneous reciprocal feedforward-feedback dynamics in the brain^40,80^. Interestingly, this interactionist perspective applies not only to perception of the visual world, but also to illusory perceptions. Identical circuitry seems to be required for both normal and illusory perceptions. Patients with Charles Bonnet syndrome exhibit activation of the visual cortex during visual hallucinations^81^. Extensive damage to the cortical visual areas results in ablation of visual imagery during sleep^82^. Patients with more circumscribed lesions exhibit more specific visual deficits both during normal wakefulness and in dreams^83,84^. Computational modeling suggests that the symmetry of visual hallucinations induced by psychedelics reflects the architecture of the primary visual cortex^85^. Unlike spontaneous activity, direct electrical activation of the visual cortex in humans leads to the perception of highly artificial phosphenes^86^. These lines of evidence together suggest that visual perception relies upon the interactions between the spontaneous dynamics and perturbations imposed onto them by the visual stimuli.

Our work suggests that the 3-6Hz traveling waves span much of the cortical surface and can, under specific circumstances, entrain firing of broadly distributed neurons. This leads to a tantalizing suggestion that this traveling cortical wave coordinates neuronal activity ultimately required for perceptual awareness. Manipulations that prevent neurons from being entrained to the traveling wave are associated with states lacking perception. Conversely, manipulations that increase the coupling of individual neurons to the traveling wave may be associated with hallucinations. The awake state is delicately balanced between these two regimes such that stimuli evoke an increase in coupling among cortical neurons which allows them to transiently form an oscillatory assembly spanning distant cortical regions.

## Methods

### Animals

All experiments in this study were approved by Institutional Animal Care and Use Committee at the University of Pennsylvania and were conducted in accordance with the National Institutes of Health guidelines. Experiments were performed on 32 adult (12–32 weeks old, 20–30 g, 21 male and 11 female) C57BL/6 mice (Jackson Laboratories). Mice were housed under a reverse 12:12 h, light: dark cycle, and were provided with food and water *ad libitum*. Inclusion criteria for mice included the following: 1) presence of visual-evoked activity in which the absolute value of the first 100 ms of post-stimulus activity exceeds five standard deviations of pre-stimulus activity and 2) presence of spontaneous activity that was not characterized as burst suppression. 25 mice were recorded from under isoflurane, ketamine and during wakefulness. 7 mice were recorded from only under isoflurane.

### Headplate implantation, habituation, and craniotomy

Headplate implantation, mouse habitation and craniotomy for mice performing isoflurane, ketamine, and awake recordings followed protocols described in Aggarwal et al, 2022^10^. Briefly, mice were anesthetized with 2.5% and maintained with 1.5% isoflurane, and secured on a stereotaxic frame (Narishege). Periosteum was exposed and bregma, lambda, and the site of the future craniotomy were marked (+1mm to −5 mm AP, +0.25 mm to +6 mm ML of bregma) on the left hemisphere. The skull was then scored and a custom designed headpiece was secured with dental cement (Metabond), cyanoacrylate adhesive (Loctite 495), and 3 skull screws (Fine Science Tools, Self tapping skull screws, 19010-10). Mice received 0.5mg cefazolin, 0.125mg meloxicam, and 7 ml of normal saline SQ post operatively, and recovered for one week. Then, mice were habituated to head fixation with body restraint with visual stimuli during one 45-minute session per day over the course of 4 days.

After completing the habituation protocol, mice were ready for recordings. Animals were anesthetized with 2.5% isoflurane in oxygen, secured onto the stereotaxic frame and maintained at 1.5% isoflurane. Before surgery, local anesthesia was injected subcutaneously in the face, scalp and neck muscles, targeting the trigeminal and occipital nerves (0.625 mg bupivacaine, 0.025%)^87^. Analgesia was supplemented with subcutaneous injection of 0.125mg meloxicam. To reduce potential swelling, 0.006mg dexamethasone was also injected subcutaneously before surgery. A 4 mm ML by 6 mm AP craniotomy was then drilled through the dental cement along the markings. A bone screw (Fine Science Tools, Self tapping skull screws, 19010-10) on the right skull bone (+2mm ML, −2mm AP) was used for a reference. A 64-electrode surface grid (E64-500-20-60, Neuronexus) was positioned over the dura (right most electrode anterior electrode at ∼ 1mm lateral and 1mm posterior to bregma).

In 14 animals, two DiI coated (Sigma-Aldrich) laminar 32 channel probes (H4, Cambridge Neurotech) were also inserted 800um into the cortex. One targeted a hole in the ECoG grid closest to V1 (−3.25 AP, −2.25 ML). The other targeted a hole in the ECoG grid closest to PPA (−1.5 AP, −1.5 ML). Both were placed using a motorized micromanipulator (NewScale Technologies). The grid and exposed dura were then covered with gel foam soaked in mineral oil to prevent desiccation. Upon completing recordings, animals were deeply anesthetized (5% isoflurane) and sacrificed. Brains were extracted and fixed in 4% paraformaldehyde (PFA) overnight prior to sectioning and histology.

### Anesthetic delivery protocol

After ECoG electrodes were positioned, animals were ready for electrophysiology recording. Mice were first given 0.6% isoflurane through a nose cone and presented visual stimuli. Isoflurane was then turned off for 30 minutes before awake recordings when visual stimuli began. After awake recordings concluded, mice were re-induced with a transient 2 minute inhalation of 2.5% isoflurane to allow for accurate administration of 100mg/kg IP ketamine. Visual stimulation and recording under ketamine anesthesia began 2 minutes after ketamine administration. Acute surgeries for animals receiving only isoflurane (n=7) were performed as described in Aggarwal, et al 2019^88^.

### Histology

Brains from the 14 animals in which laminar probes were inserted, were sectioned at 80µm on a vibratome (Leica Microsystems). Sections were mounted with a medium containing a DAPI counterstain (Vector Laboratories) and imaged with an epifluorescence microscopy (Olympus BX41) at 4x magnification. The stereotaxic locations of the laminar probes were determined using post-mortem histology by comparison to the brain atlas^89^.

### Visual Stimulation

Two sets of visual stimuli were used in the study. The first consisted of 100 trials of 10 ms flashes of a green LED (650 cd/m^2^), covering 100% of the mouse’s right visual field, delivered at a random interstimulus interval drawn from a uniform distribution between 3 to 4 seconds. In 17 animals, 240 trials of 100ms full flashes of a CRT monitor (Dell M770, refresh rate 60 Hz, maximum luminance 75 cd/m^2^), placed 23 cm away at an angle of 60% from the mouse’s nose, thereby covering 70% of the mouse’s right field of view. The monitor full field flashes varied in luminance’s (2%, 11%, 25%, 44%, 75%, 100%) in random order at a random interstimulus time interval between 3 and 5 seconds. The corresponding luminance values are shown in Figure 6.

### Electrode registration

Histological localization laminar probes was used to align the ECoG grid to stereotaxic coordinates. The angle of the line formed by two laminar probe insertion sites and the AP axis was defined as θ. Each electrode on the ECoG grid was then assigned a location based on its Euclidean distance from the two laminar probe sites. Finally, the resultant grid location matrix was multiplied by a rotation matrix (*R*) to obtain the final electrode positions in stereotaxic coordinates.

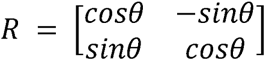

The resulting ECoG coordinates were then compared to photographs of the ECoG grid relative to bregma and lambda to verify coordinate assignment.

### Electrophysiology and preprocessing

In 6 mice, signals were amplified via a Neuralynx headstage (HS36), digitized with a Cheetah 64 acquisition system (Neuralynx, ERP-27, Lynx-8), and collected at a rate of 3030.3Hz/channel. In the remaining mice, signals were amplified and digitized on an Intan headstage (Intan, RHD2132) connected to an Omniplex acquisition system (Plexon, Omniplex), and collected a sampling rate of 40KHz/channel.

The LFP data were first downsampled to 1 kHz and then filtered between 0.1 Hz and 325 Hz using *firls.m* and *filtfilt.m* functions in MATLAB, to minimize phase distortions. Channels that contained line noise and trials with excessive movement artifacts were eliminated manually. To minimize the impact of volume conduction, the average signal across all electrodes was subtracted from the LFP. All subsequent analyses were conducted using custom-made MATLAB code unless otherwise specified.

### Selection of V1 ECoG electrode

In 18 mice, laminar probes were not inserted and therefore grid electrodes were not aligned to stereotaxic coordinates. In these mice, the V1 electrode was identified neurophysiologically ^10,88^. The latency of onset of the visual-evoked potential was calculated for each ECoG electrode as the time point at which the post-stimulus average LFP exceeded 3 standard deviations above the pre-stimulus baseline for 3 consecutive time points. The electrode which had the lowest latency of onset was defined as V1.

In the 14 mice in which laminar probes were inserted and the stereotaxic positions of the ECoG probes were inferred, the electrode closest to stereotaxic V1 (−3.25 AP, −2.25 ML) was chosen for each animal. To confirm that the chosen electrode neurophysiologically corresponded to V1, the latency of onset of the VEP at each grid electrode was also computed as above. The position of the stereotaxically identified V1 electrodes was within 1-2 electrodes (<1000um) in distance and the onset was within 2 ms of the electrode with the earliest latency of onset in all mice.

### Current Source Density Analysis (CSD)

LFP from the laminar probes was converted in CSD by computing the second spatial derivative ^90^:

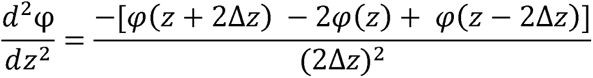

where φ is the LFP, z is the vertical coordinate depth of the probe, and Δz is the interelectrode distance (25 µm). Estimation of the CSD at the boundary electrodes was projected using the Vaknin estimation procedure^91^. Channels with the earliest current sink in the V1 probe were assigned as layer 4 (granular layer). Subsequent sinks were found above and below layer 4 in layers 2/3 and layer 5 of the V1 probe. The channels of the PPA probe were assigned a laminar structure based on their distance from the cortical surface as described in Aggawral et. al 2022^10^. Laminar data was only included in analysis if the channels could clearly be assigned to layers in this manner (11 out of 14 mice).

### Wavelet analysis

Continuous wavelet transform with Morlet wavelets implemented in *contwt.m* (0.1 Hz to 150 Hz, with a step-width 0.25 Hz and normalized amplitude) was used to calculate the power, phase, and frequency characteristics of LFP or CSD (http://paos.colorado.edu/research/wavelets/) ^92^.

### Inter-trial Phase Coherence (ITPC) Analysis

Intertrial phase coherence (ITPC) is a measure of phase consistency of LFP filtered at specific frequency bands. The phase of the LFP in each frequency band was extracted using continuous wavelet transform. The ITPC was computed as the circular across trial average of such unit length phase vectors^93^.

### Filtering data for wave analysis

LFP or CSD data was filtered into (3-6Hz) using the inverse wavelet transform, *invcwt.m*, (available at: http://paos.colorado.edu/research/wavelets/) ^92^. All wavelet coefficients outside the desired frequency band were set to zero. As we have shown previously^10^ the results are largely unaffected by the choice of the filtering method.

### Construction of Analytical 3-6 Hz Signals

Hilbert transform was used to extract the analytical signal of LFPs or CSDs filtered for 3-6Hz frequencies. This resulted in series of complex numbers, in which the modulus of the analytical signal corresponds to the instantaneous amplitude and the arctan of the analytical signal corresponds to the instantaneous phase.

### Complex Singular Value Decomposition (SVD)

Detailed methods and examples describing complex SVD analysis of filtered LFP can be found in Aggarwal et. al^10^. Briefly, analytical signal (spanning from 1600ms to 100 ms of pre-stimulus activity for spontaneous trials 500ms of pre-stimulus to 1000 ms of post-stimulus activity for stimulus evoked trials) from ECoG electrodes formed matrix **A** for each trial. Oscillatory modes were extracted from A by performing singular value decomposition, which factorizes A into mutually orthogonal modes:

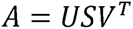

the spatial and temporal components of each mode, respectively. The diagonal real-valued S contains singular values (*λ’s*) and corresponds to the fraction of the total variance explained by each mode.

The spatial amplitude and spatial phase of the *i-th* mode is the modulus and the arctan of *U_(*,i)_* * *λ_i_*. Temporal amplitude and temporal phase are the modulus and the arctan of *V_(*,i)_* * *λ_i_* respectively.

### Reliability of visually responsive modes

The temporal amplitudes of the first ten singular modes were computed for each single trial, as above, and then normalized to their mean and standard deviation over 400 ms of pre-stimulus activity. Modes in which the temporal amplitude exceeded 6 standard deviation above pre-stimulus activity within the post-stimulus window (1000ms post-stimulus for 3-6Hz activity), were defined as visually responsive modes.

### Defining the most visually responsive mode

The mode that displayed the greatest increase in temporal amplitude during the post-stimulus period compared to pre-stimulus activity, was defined as the most visually responsive mode.

### Consistency of Spatial Amplitude Loading using Earth Movers Distance

Earth Movers Distance was calculated for the spatial amplitude of the most visually evoked modes for each pair of trials using the Matlab function *emd.m*. This was done for each mouse in each experimental condition. Next, the probability density function (PDF) of the Earth Movers Distance was calculated. 95% confidence intervals obtained through bootstrapping.

### Consistency of Spatial Amplitude Loading using Cosine Similarity

To determine the similarity in the spatial activation of the most visually evoked modes across trials, the cosine distance of the spatial amplitude of the most visually evoked mode was computed between pairs of single trials, within each mouse in each condition. Subsequently, the PDF of the cosine similarity (1-cosine distance) was computed for all pairs of trials across all mice for each state. The cosine similarity of spatial amplitudes in mice under anesthesia is subtracted from the PDF of the cosine similarity of spatial amplitudes in awake mice.

### Spatial phase offset from V1

For each single trial, the phase offset from V1 was computed by extracting the spatial phase of the most visually responsive mode at each electrode and projecting it onto a unit circle. To find the average phase angle and consistency of each channel’s phase offset relative to V1, the circular mean was computed across trials within each animal^94^. The direction of the mean vector corresponds to the angle of phase offset whereas the magnitude corresponds to 1-variance.

In the 14 animals in which ECoG electrode stereotaxic positions were identified, the circular mean and variance were computed across trials and animals at each stereotaxic location.

### Spatial phase gradient

Using *phase_gradient_complex_multiplication.m*, written by Lyle Muller (available at: https://github.com/mullerlab/wave-matlab), the spatial phase gradient was calculated on a single trial basis by iteratively multiplying the complex valued spatial component of the most sensory evoked mode at one electrode location to the complex conjugate of the spatial component of the next electrode in the medial-lateral and anterior-posterior directions^9^. The average gradient over trials was calculated by projecting each trial’s gradient vector at each position onto a unit circle and computing the circular mean. The angle of the resultant vector corresponds to the direction of spatial phase advancement, whereas the magnitude of the resultant vector corresponds to 1-variance in the direction of the gradient over trials.

### Averaging stereotaxic coordinates over animals

A query set of stereotaxic locations was defined from −5 to 0 mm AP and −3.5 to 0 mm ML, with 0.5mm spacing. The weight of each electrode at each query site was assumed to be Gaussian function of the distance between the electrode and the query location:

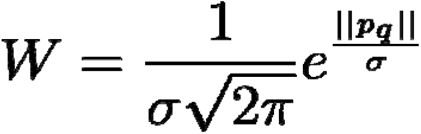

Where p_q_ is the Euclidean distance between the electrode p and query location q, σ= 0.15 is the standard deviation. The weight of all electrodes outside of 0.3 mm radius of the query position was set to zero.

### Spike Sorting

The same 11 mice that were used for laminar analysis also had their probe data analyzed to identify individual neurons. Kilosort^95^ was used for spike sorting, and then the resulting spikes were visually checked for accurate waveform clustering using Phy. Neurons with a firing rate below 0.5 spikes per second were removed from the analysis.

### Spike Field Coherence (SFC)

Was computed using the method by Tort et al^96^. CSD (filtered at 3-6Hz) from the same electrode as the action potential was used for phase.

### Time-Warped Raster Generation

For each neuron, time-warped raster plots were created with three cycles of 3-6Hz filtred CSD from each trial. A horizontal stretch and shift value was found for every trial’s CSD such that its correlation with the first trial is maximized. Optimization was implemented in *fminsearch.m* function (Matlab 2020b, Mathworks). Once the optimal stretch and shift values were obtained for each trial, the transformations were applied to the trial’s corresponding spike times.

### Spike Coherence

Coherence between spike trains of each pair of V1 and PPA neurons within the same mouse were calculated on a single trial basis using the Matlab function *wcoherence.m,* (Matlab 2020b, Mathworks). Spike coherence was then averaged over all trials for each pair of neurons.

### Statistical Analysis

In all cases, the threshold for statistical significance was set at *p<0.05.* To determine the statistical significance of the ITPC, the observed ITPC were compared to time shifted surrogates (n = 100 sets, 100 trials per set), using a one tailed t-test.

To determine the effects of brain state on the wave distributions, reliability of visually evoked modes, SVD rank of most visually evoked mode, consistency in phase offset from V1 of most visually responsive mode across luminance, and spike coherence, a Kruskal Wallis test was first performed. If there was a statistically significant effect of brain state, pairwise hypotheses were tested using Dunn-Sidak *post hoc* analysis, thus correcting for multiple comparisons.

To determine if the proportion of neurons entrained to the phase of the slow wave was different across brain states, a Chi Square test for Homogeneity was preformed. Pairwise hypotheses were tested using the *post hoc* Tukey’s Honest Significant Differences Test among Proportions, thus correcting for multiple comparisons.

To assess the significance of the spatial phase and gradients at each location, a Rayleigh test was conducted to determine whether there was a deviation from circular uniformity. The p-values resulting from this test were then corrected for multiple comparisons using the Bonferroni method (64 ECoG channels within individual mice and 77 query stereotaxic locations across mice).

### Presentation of previously printed figures

Awake data in Figures 6b is adapted from Sup. Figure 8b Aggarwal et. al, 2022^10^.

## Supporting information

SupFigures

## 2 Author Contributions

Conceptualization: AA, DC, MBK, and AP

Data curation: AA, HC, DC, and AP

Formal analysis: AA, JL, and AP

Funding acquisition: AA, DC, MBK, and AP

Writing – original draft: AA, MBK, and AP

Writing – review & editing: AA, JL, MBK, DC, and AP

## Acknowledgments

We also want to thank Dr. Joseph Cichon, Dr. Andrew McKinstry-Wu, Connor Brennan, Andrzej Z. Wasilczuk, Claudia Heymach and Ethan Blackwood for helpful discussions. This research was supported through the Translational Neuroscience Initiative from the Penn Medicine Translational Neuroscience Center (PMTNC), RO1 GM088156 (MBK), RO1 GM124023 (AP), T32 EY007035 (AA), F30 EY029931-01A1 (AA).

## 4 Supplementary Material

6 supplementary figures are submitted.

## 5 Data Availability Statement

The raw data and MATLAB code presented in this manuscript will be made available upon request.

