## Supplementary material for "Neural assemblies coordinated by cortical waves are associated with waking and hallucinatory brain states": SupFigures

### 1.1 Supplementary Figures

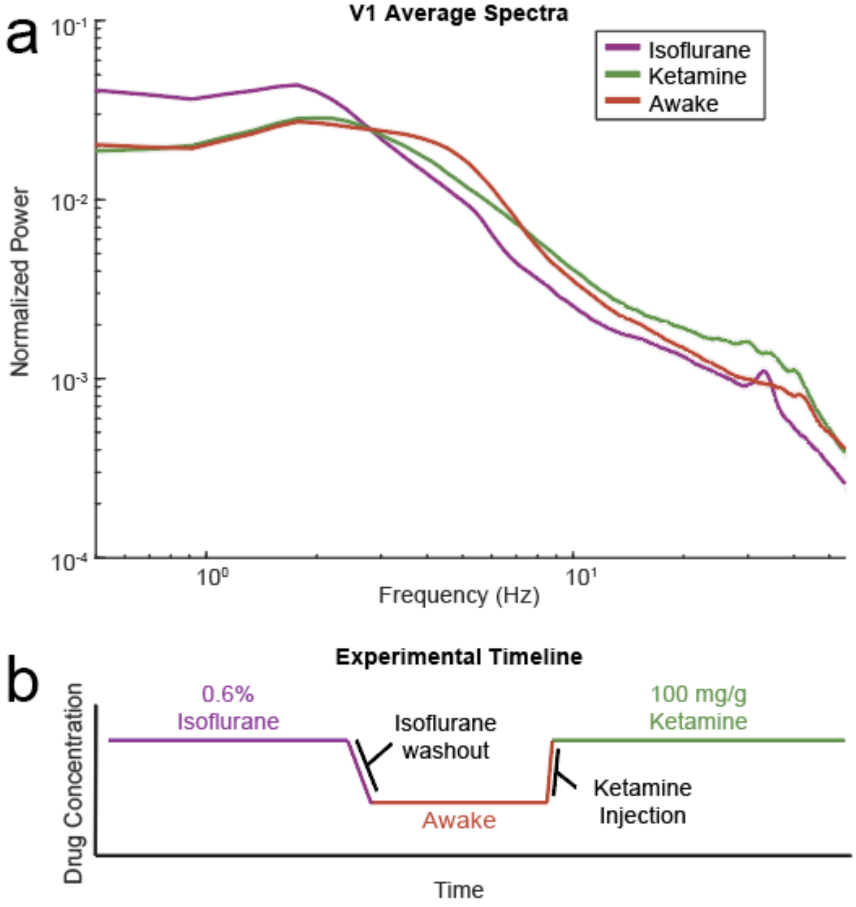


**Supplementary Figure 1: Baseline spectrum and experimental timeline**

A. Average power spectrum of spontaneous activity recorded from an ECoG on top of V1 for awake mice (red) and mice under isoflurane (purple), and ketamine (green). Shading indicates the 95% confidence intervals. LFP recorded from awake mice and mice under ketamine have higher power at higher frequencies whereas LFP recorded from mice under isoflurane have a predominance of low frequency oscillations.

B. Experimental timeline (methods). Mice were first given 0.6% isoflurane through a nose cone, then given 30 mins for isoflurane washout before awake recordings began, then given 100 $\mu$g/g ketamine IP

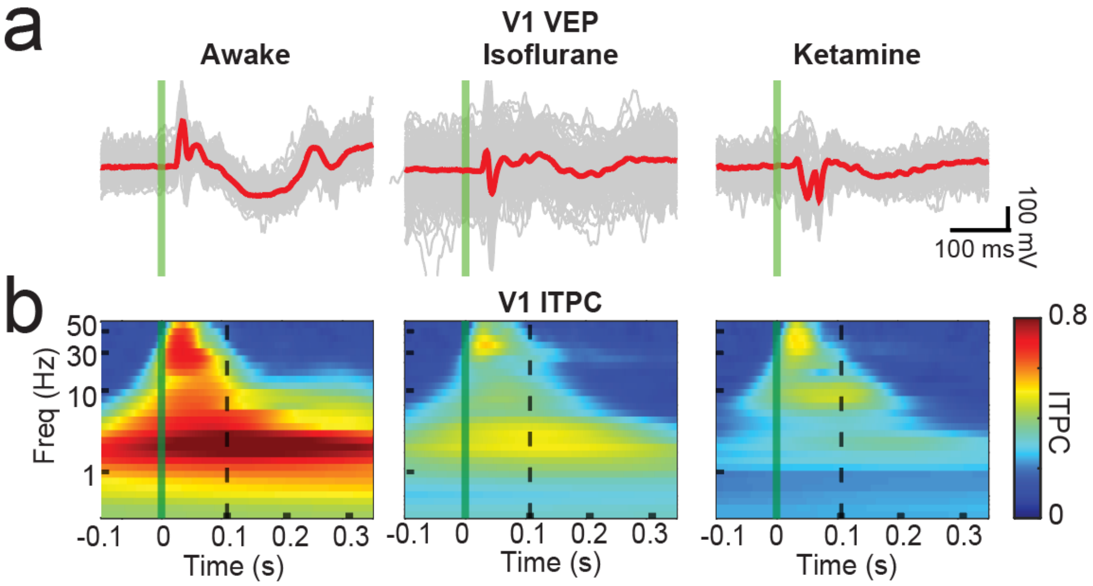


**Supplementary Figure 2: Visual stimuli of sufficient intensity elicit gamma coherence in V1 in all three states**

A. Examples VEPs in V1 elicited by an LED flash in all three states: awake (left), isoflurane (middle), ketamine (right). Thin grey lines denote single trials, thick red lines denote average VEPs, and the stimulus occurs at the green vertical line.

B. Intertrial phase coherence averaged across animals in all three states elicited by an LED flash: awake (left), isoflurane (middle), ketamine (right). Time is on the x-axis, frequency is on the y-axis, and color denotes ITPC. The stimulus occurs at the green vertical line and 100ms post-stimulus occurs at the dashed black line.

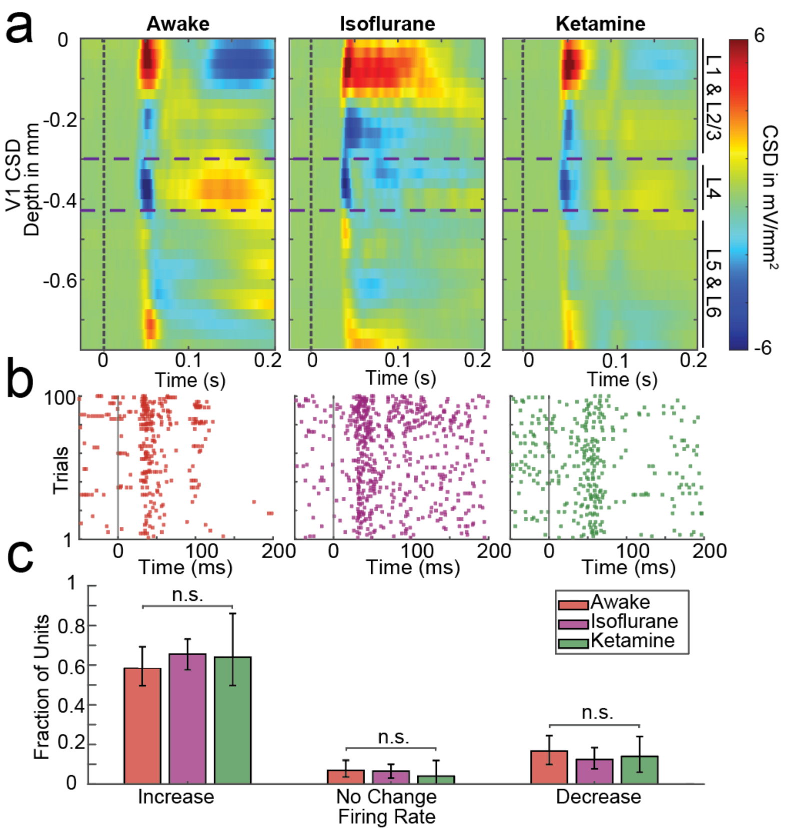


**Supplementary Figure 3: V1 receives thalamic input and is activated by stimuli in all three brain states**

A. Current source density (CSD) averaged over mice from laminar electrode arrays placed in V1 in animals that are awake (left), under isoflurane (middle), or under ketamine (right). Current sinks are depicted in blue and current sources are shown in red. The purple dashed lines indicate the granular layer boundaries (L4). Note that the early VEP is preserved; however, later aspects of the visual evoked response are diminished in amplitude under isoflurane and ketamine

B. Raster plot of an example neuron in V1 in each state, with time on the x-axis, trial on the y-axis and each dot corresponding to the time of an action potential. Note that these cells increase in firing to the stimulus in all three brain states.

C. Fraction of units that increase (right), do not change (middle) or decrease (left) in firing rate after stimulus presentation. Note that the composition of neural responsiveness does not change with changing brain state.

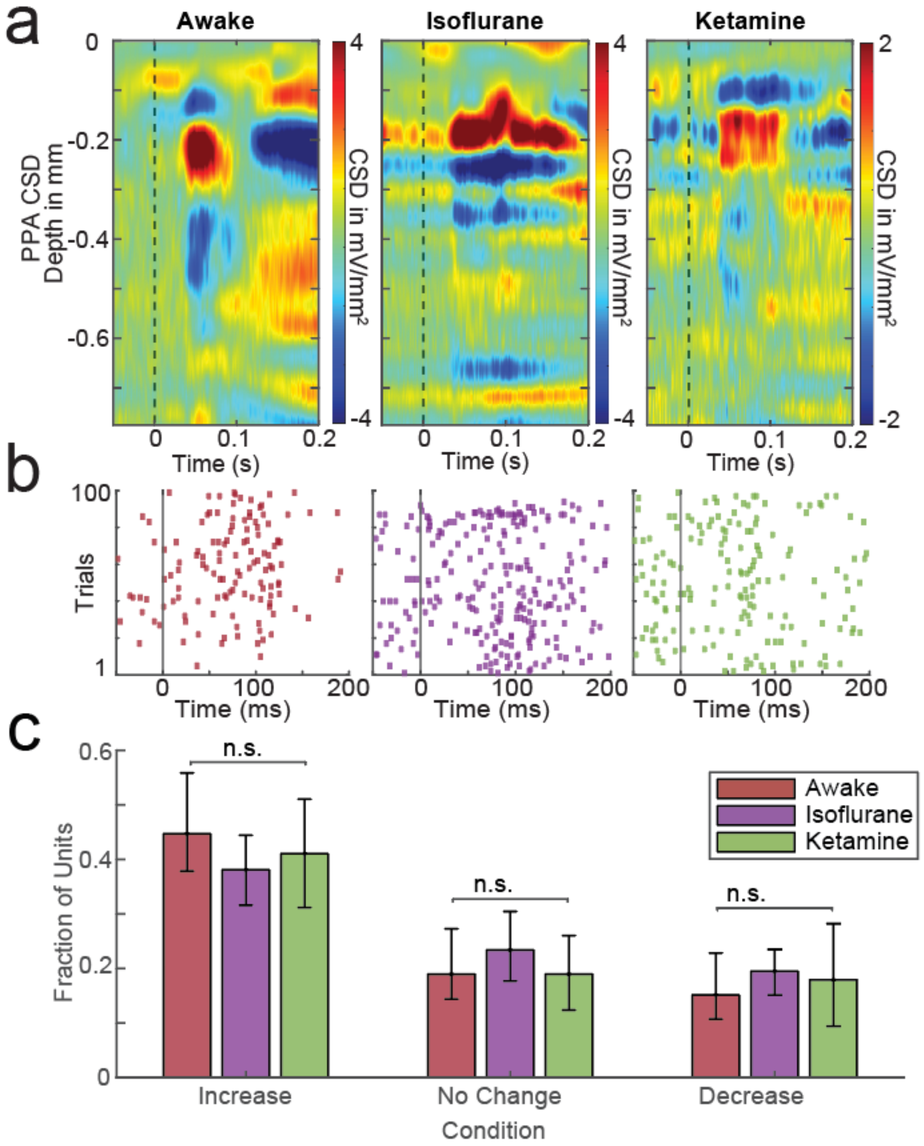


**Supplementary Figure 4: PPA receives input and is activated by stimuli in all three brain states**

D. PPA CSD averaged over trials in a representative mouse during wakefulness (left), under isoflurane (middle), or under ketamine (right). Current sinks are depicted in blue and current sources are shown in red.

E. Raster plot of an example neuron in PPA in each state, with time on the x-axis (the black vertical line denotes the flash), trial on the y-axis and each dot corresponding to the time of an action potential. Note that these cells increase in firing to the stimulus in all three brain states.

F. Fraction of units that increase (right), do not change (middle) or decrease (left) in firing rate after stimulus presentation. Note that the composition of neural responsiveness does not change with changing brain state.

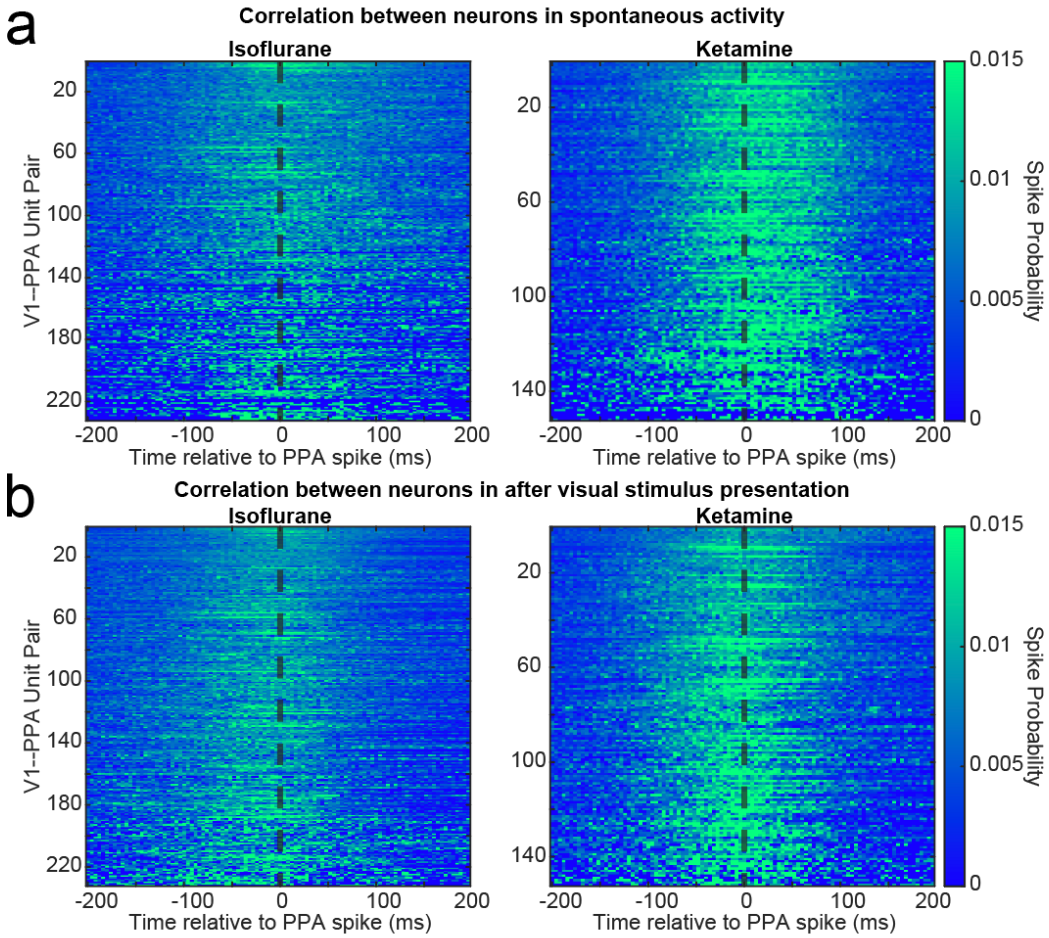


**Supplementary Figure 5: Correlations between V1 and PPA neurons do not change significantly after stimulus presentation when animals are under isoflurane or ketamine**

A. Cross-correlograms between PPA and V1 neurons entrained by the 3-6 Hz wave during the 500 ms before visual stimulation in animals under isoflurane (left) and under ketamine (right). Each row is an individual PPA V1 pair. Probability of firing is shown by color.

B. Same as in A but for 500 ms after the stimulus in animals under isoflurane (left) and under ketamine (right).

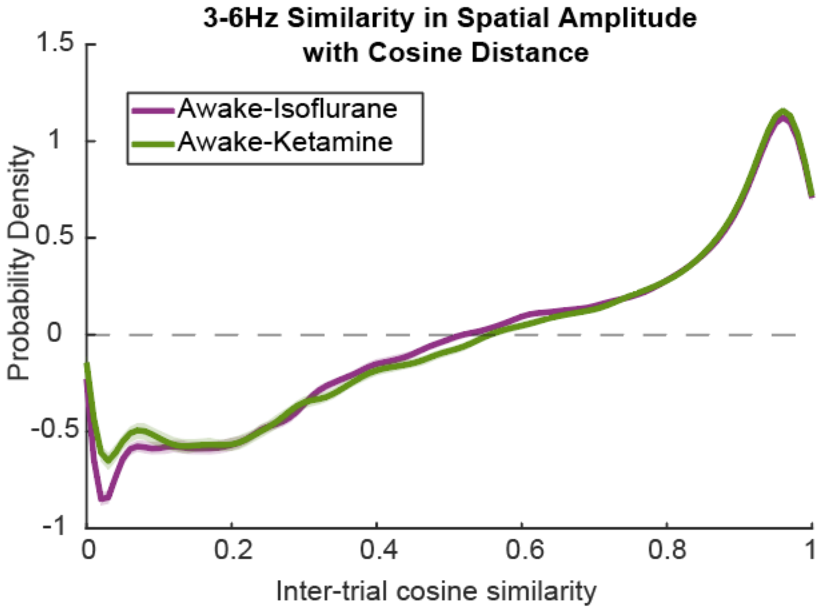


**Supplementary Figure 6: Cosine similarity of spatial amplitude of single trials is higher in animals that are awake compared to when animals are under ketamine or isoflurane**

The difference in the cosine similarity between the 3-6Hz spatial amplitude of the most visually responsive SVD mode between awake and isoflurane (purple) and awake and ketamine (green). The dashed black horizontal line denotes no difference in the cosine distances. Note that when mice are awake, the spatial activation across electrodes is more similar from trial to trial than when animals are under anesthesia.

116
